# ApeA cleaves genomic RNA to defend against RNA phage infection

**DOI:** 10.64898/2026.03.20.713152

**Authors:** Arina Drobysheva, Manuel Velasco Gomariz, Shazeb Ahmad, Sarah Reichardt, Jens Hör

## Abstract

To protect themselves against bacteriophage infection, bacteria encode a vast diversity of antiphage defense systems. However, the mechanisms of action of most of these systems have exclusively been studied using phages with DNA genomes as the models, while phages with RNA genomes remain understudied. Here, we investigate how the defense system ApeA confers resistance against RNA phage infection. We show that two ApeA homologs, Ec1ApeA and Ps2ApeA, protect against a variety of single-stranded RNA phages. Focusing on Ec1ApeA, we find that it senses infection through a conserved pocket that likely binds an RNA structure in the phage genome. This activates the HEPN (higher eukaryotes and prokaryotes nucleotide-binding) RNase domain of Ec1ApeA which consequently cleaves the phage genomic RNA to restrict replication. In contrast to many other described defense systems, Ec1ApeA activity directly stops viral replication without inducing cell death, establishing ApeA as a non-abortive defense system that protects against RNA phages. Our results add to the increasingly diverse targets of antiviral HEPN RNases and provide insights into the understudied field of RNA phage defense.

## Introduction

Bacteria employ a plethora of antiviral strategies to defend against bacteriophage predation. For example, bacteria can modify the proteins that phages use as receptors to limit adsorption and therefore infection^1^. In addition, bacteria employ a vast variety of active antiphage defense systems to block phage replication, which together constitute the bacterial immune system^2–5^. Recent bioinformatics studies with follow-up validation experiments have identified a staggering number of defense systems that are active against tailed double-stranded DNA phages of the class *Caudoviricetes*^6–8^. This revealed that bacterial immunity is much more complex and diverse than previously anticipated^9^. Importantly, many of these defense systems were shown to be the evolutionary origins of innate immune systems of eukaryotes, thus unveiling that many antiviral strategies are conserved across the tree of life^10–12^.

While it is well known that eukaryotes use RNA sensors such as RIG-I or MDA5 to detect infection by viruses with RNA genomes^13^, little is known about RNA sensors that bacteria might use to defend themselves against infection by phages with RNA genomes. RNA phages are non-tailed, replicate without a DNA intermediate, and were historically considered to be rare in nature^14^. Recent metatranscriptomics studies have overturned that assumption, revealing that RNA phages are widespread across diverse ecosystems, in some of which RNA phages constitute the majority of detectable RNA viruses^15,16^. Among cultivated RNA phages, virulent members of the family *Fiersviridae* (formerly *Leviviridae*), such as the model phages MS2 and Qβ, are the best understood. They possess highly compact ∼3500–4000 nt positive-sense single-stranded RNA (+ssRNA) genomes and encode only four proteins^17,18^. Despite their minimal genomes, fiersviruses and related members within the class *Leviviricetes* are the most abundant RNA phages in nature^15,16^, showcasing their evolutionary success and the selective pressure they likely impose on bacterial populations.

In stark contrast to immunity against tailed DNA phages, only a handful of defense systems have been shown to protect against RNA phages. The best-understood of these systems are adaptive type III CRISPR-Cas systems that can be programmed to restrict infection by model RNA phages^19–21^. Consistently, metagenomic studies of type III CRISPR-Cas systems with a reverse transcriptase domain in their acquisition machinery have identified spacers acquired from RNA phage genomes^15,22,23^. This suggests that these systems also defend against RNA phages in nature. In addition to adaptive CRISPR-Cas systems, a few innate systems have been shown to stop RNA phage replication. Examples include bNACHT (bacterial NACHT module-containing) proteins^24,25^, type I Zoria^26,27^, Ataecina^28^, and Candamuis^28^. Yet, only RNA phage defense by bNACHT proteins has been functionally characterized^25^, highlighting that there is a large gap in our understanding of how bacteria sense and stop RNA phage infection.

In this study, we show that the ApeA defense system protects against RNA phages of the *Fiersviridae* family. ApeA was identified in a screen for defense systems, where it was found to protect against several DNA phages^7^. In comparison to most other defense systems, ApeA consists of only a single protein that comprises an N-terminal domain of unknown function as well as a C-terminal HEPN (higher eukaryotes and prokaryotes nucleotide-binding) RNase domain (Figure 1A). HEPN domains are widespread across the tree of life and are involved in diverse functions, for instance as the effector nuclease in antiviral systems^29,30^. To name a few examples, in mammals, RNase L contains a HEPN domain^31^, while in bacteria, several CRISPR-Cas-related proteins such as Cas13^32^, Csm6^33^, and Csx1^34^, as well as several toxin-antitoxin systems^35–38^ contain HEPN domains. We show that ApeA senses an RNA structure in the genomic RNA of the Qβ-like RNA phage FrSangria via a conserved pocket, which subsequently activates the HEPN domain to cleave the phage genome. Thus, ApeA stops RNA phage replication via direct genome degradation. Our study reveals a new mechanism in bacterial immunity and sheds light on how bacteria defend themselves against the understudied class of RNA phages.

**Figure 1.**
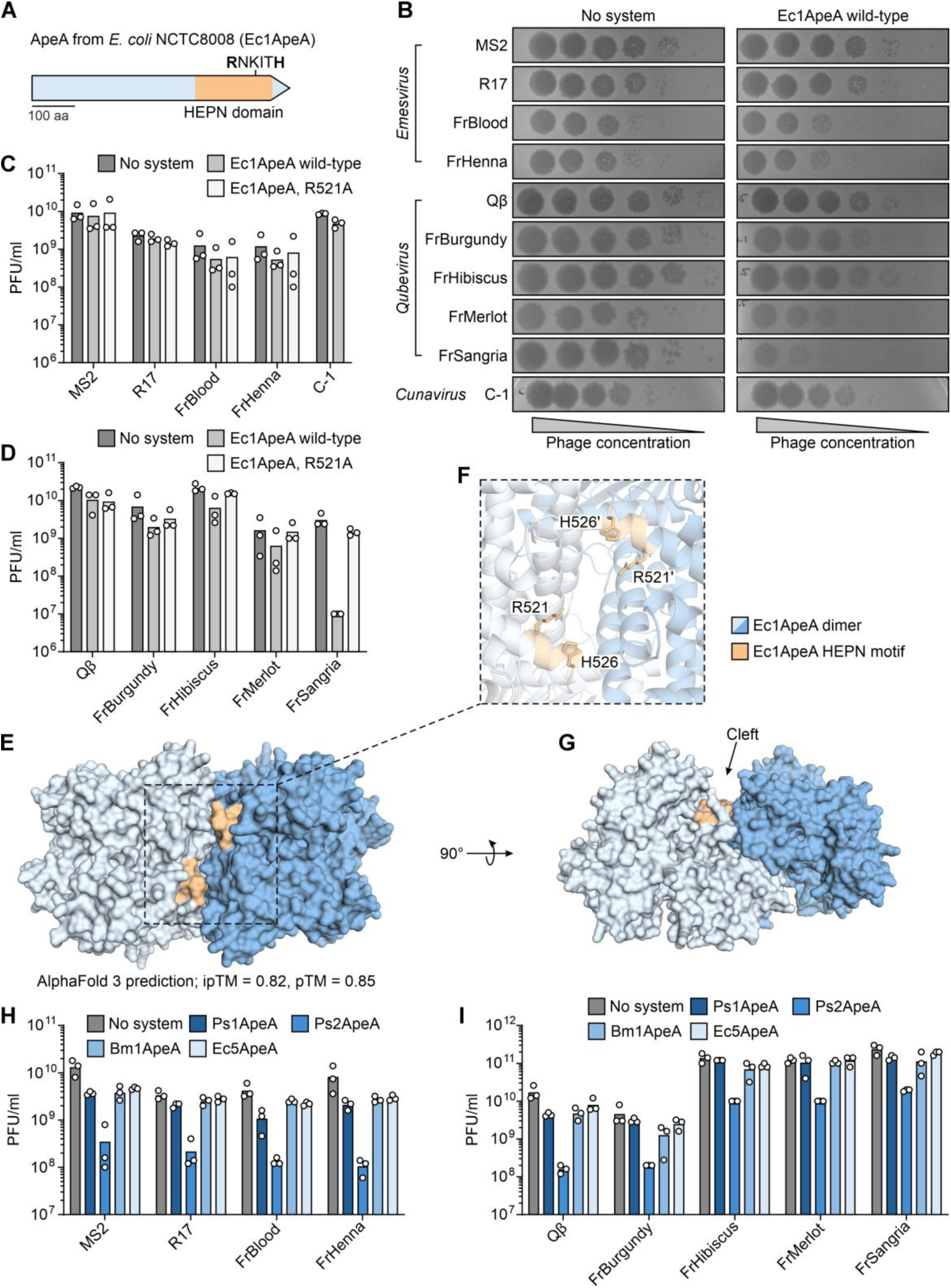
ApeA confers resistance against RNA phage infection. **A,** Schematic representation of the ApeA protein from *E. coli* NCTC8008 (Ec1ApeA). The 601 amino acid (aa) long protein comprises a C-terminal HEPN RNase domain with the active site motif RNKITH. **B,** Plaque assays showing the defense phenotype of wild-type Ec1ApeA against RNA phages in comparison to the control. Data are representative of three biological replicates. **C, D,** PFU quantification of plaque assays shown in **B** as well as the Ec1ApeA R521A mutant. Bars represent the average of three biological replicates with individual data points overlaid. **E,** Structure of the Ec1ApeA homodimer, predicted by AlphaFold 3^42^. ipTM = 0.82, pTM = 0.85. Plot of the predicted alignment error is shown in **Fig. S1B**. **F,** Zoom-in of the HEPN motifs of the two Ec1ApeA monomers. **G,** The HEPN motifs of the Ec1ApeA monomers form the base of a deep cleft. **H, I,** PFU quantification of RNA phages infecting *E. coli* expressing different ApeA homologs. Bars represent the average of three biological replicates with individual data points overlaid.

## Results

### ApeA is a HEPN domain-containing protein that protects against RNA phages

ApeA was originally identified in a large screen for defense systems and consists of a single gene encoding a ∼600-aa protein with an N-terminal domain of unknown function as well as a C-terminal HEPN RNase domain^7^ (Figure 1A). ApeA is widespread in bacteria, such as in the Bacillati, the FCB and PVC superphyla, the Pseudomonadota, and the Spirochaetota^7^. When expressed under its native promoter on a medium-copy plasmid in *E. coli*, ApeA from *Escherichia coli* NCTC8008 was shown to strongly protect against the DNA phage T2 and to weakly protect against the DNA phages T3, T4, T5, T7, and φV-1^7^. In a recent preprint by others, this ApeA homolog was renamed Ec1ApeA and was shown to protect against a variety of additional DNA phages^39^. Motivated by the HEPN RNase domain of ApeA, we hypothesized that it might also protect against phages with RNA genomes and thus set out to study the defense potential of Ec1ApeA against a diverse panel of ssRNA phages. As shown before^7^, we did not observe defense against the model RNA phages MS2 and Qβ (Figure 1B–D). By contrast, Ec1ApeA conferred strong protection (∼300-fold reduction in PFUs (plaque-forming units)) against FrSangria, a phage related to Qβ in the genus *Qubevirus*^40^ (Figure 1B, D). Additionally, presence of Ec1ApeA substantially reduced the plaque size of the RNA phages FrBurgundy, FrHibiscus, and FrMerlot, all of which are close relatives of FrSangria (>90% sequence identity^25^) (Figure 1B). This suggests that these *Qubevirus* phages harbor unique features that allow Ec1ApeA to sense and stop the infection. As expected^7^, we also observed strong defense against the DNA phages T2, T4, and T5, which are entirely unrelated to our panel of RNA phages (Figure S1A).

For RNase activity, two HEPN domains must dimerize, which can occur either intramolecularly, such as in Cas13^41^, or intermolecularly, such as in HepT of the HepT/MntA toxin-antitoxin system^37^. Like HepT, ApeA only has one HEPN domain encompassing the conserved active site motif Rx4-6H, which is RNKITH in the case of Ec1ApeA (Figure 1A). As expected, structural analysis with AlphaFold 3^42^ predicted a high-confidence interaction between two Ec1ApeA monomers, suggesting that Ec1ApeA forms a homodimer (Figure 1E, S1B). The two HEPN motifs were placed in close proximity to each other to form the base of a deep cleft (Figure 1F, G). This cleft is a typical feature of HEPN nucleases and is usually involved in the alignment of the RNA substrate^30^. A single amino acid substitution (R521A) in the HEPN motif of Ec1ApeA fully abrogated defense against FrSangria and the tested DNA phages, indicating that RNase activity of ApeA is required for defense (Figure 1C, D, S1A).

To understand whether RNA phage defense is a common property of ApeA proteins, we then conducted a search for additional ApeA homologs using PSI-BLAST^43^ and Foldseek^44^ and built a phylogenetic tree of the identified homologs (Figure S1C, Table S1). Based on this tree, we selected 4 ApeA homologs from diverse clades with ∼21–31% amino acid identity compared to Ec1ApeA. Additionally, we selected a distant 467-aa minimal homolog from *Brucella pseudomallei*, identified through a phmmer^45^ search, which had no detectable sequence similarity based on BLAST analysis (Figure S1C). We named the chosen proteins Bm1ApeA, Bp1ApeA, Ec5ApeA, Ps1ApeA, and Ps2ApeA (Figure S1C, Table S1). Synthetic DNA encoding the proteins was cloned into a medium-copy plasmid under the control of an IPTG-inducible promoter and transformed into *E. coli*. Next, we tested the defensive potential of these proteins against our panel of RNA phages as well as several DNA phages. Ps2ApeA showed robust defense against all tested RNA phages from the *Emesvirus* and *Qubevirus* genera (Figure 1 H, I) as well as the DNA phages P1 and T2 (Figure S1D). By contrast, Bm1ApeA, Ec5ApeA, and Ps1ApeA only protected against various DNA phages (Figure S1D). We were not able to transform and test Bp1ApeA, presumably due to toxicity in our model host. Overall, these results demonstrate that ApeA does not solely protect against DNA phages, but instead provides broad defense against both RNA and DNA phages.

### ApeA cleaves the phage genome to stop the infection

Since Ec1ApeA strongly protected against the RNA phage FrSangria when expressed from a medium-copy plasmid under the control of its native promoter without the need to overexpress it (Figure 1B, D), we focused on the mode of action of this ApeA homolog. Typically, defense systems stop phage replication either directly, e.g. by cleaving the phage genome such as in the case of restriction-modification systems^46^, or indirectly, e.g. by inducing cell death^47^, growth arrest^48^, or modifying the emerging virions^49^. To assess whether ApeA operates directly or indirectly, we first monitored bacterial growth of cells expressing Ec1ApeA following infection with FrSangria in liquid culture and compared it to control cells expressing RFP (red fluorescent protein) (Figure 2A). At low MOI (multiplicity of infection), Ec1ApeA fully protected the culture against the infecting RNA phage. At high MOI, expression of Ec1ApeA enabled continuous growth of the culture, albeit at a slower rate than the uninfected control (Figure 2A). Importantly, no culture collapse was observed, as is typically the case for abortive infection systems that induce cell death before the infecting phage can complete replication^50,51^. The mutation in the HEPN domain mirrored the results of the control, as expected (Figure 2A). In contrast to these observations, high MOI infection of bacteria expressing Ec1ApeA with the DNA phage T5 caused full growth arrest that was not overcome within 8 h post infection, suggesting a potential difference in mechanism between RNA and DNA phages (Figure S2A). Neither for RNA nor for DNA phages induced cell lysis could be observed, demonstrating that ApeA does not induce cell death (Figure 2A, S2A).

**Figure 2.**
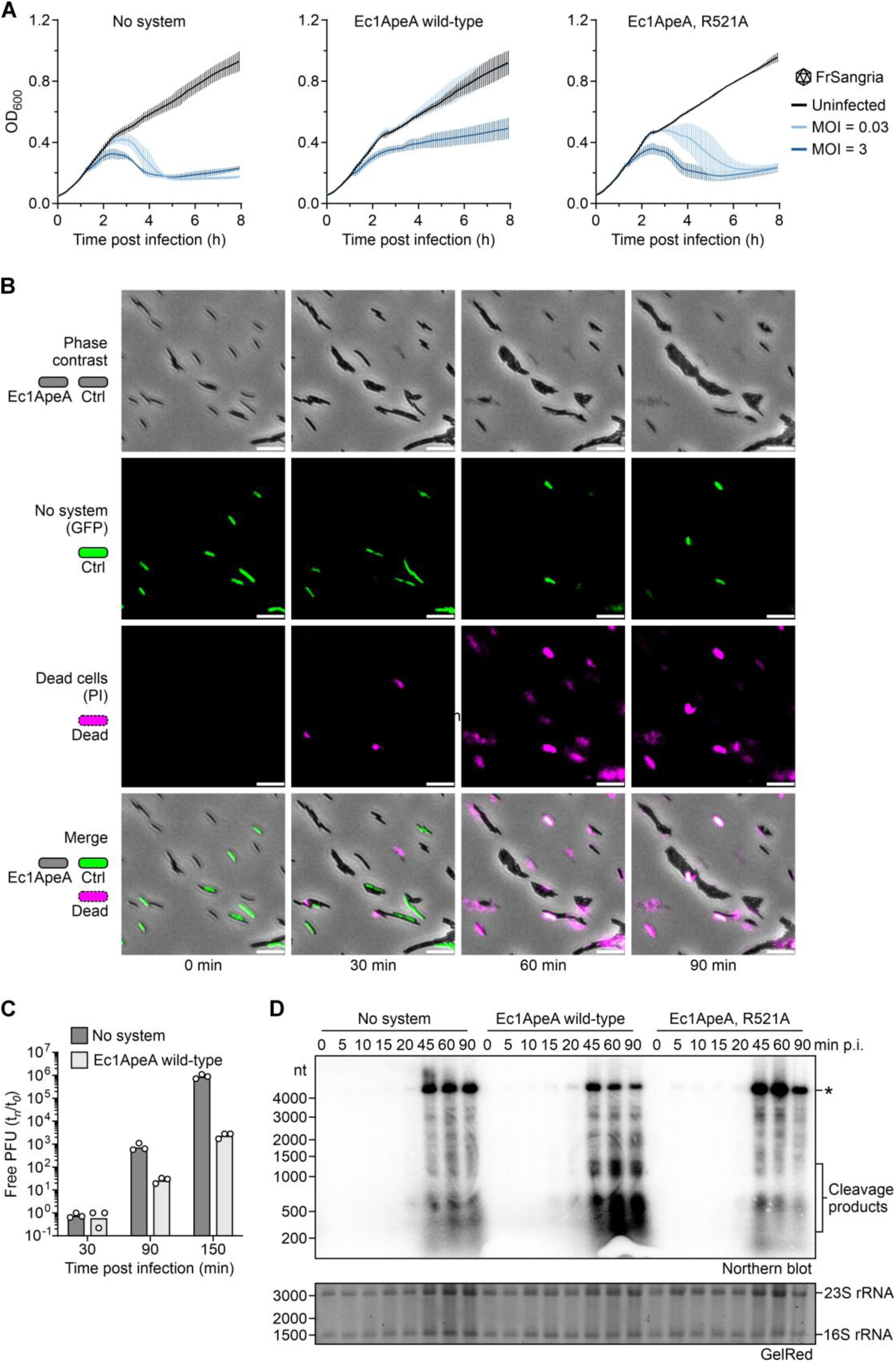
ApeA inhibits RNA phage replication by cleaving the phage genome. **A,** Growth curves of *E. coli* expressing RFP (no system), wild-type Ec1ApeA, or the Ec1ApeA R521A mutant, either uninfected or infected with FrSangria at an MOI of 0.03 or 3. The average with s.d. of three biological replicates is shown. **B,** Time-lapse microscopy of live *E. coli* cells expressing GFP (green) mixed with cells expressing Ec1ApeA (black). Cells were infected with FrSangria at an MOI of 5 in the presence of propidium iodide (PI, magenta) and incubated at 37°C in an agar pad. The three channels (phase contrast, GFP, PI) are presented separately as well as merged. Scale bar, 10 µm. Data are representative of three biological replicates. **C,** Phage replication assay. PFU quantification of FrSangria from the supernatant of *E. coli* expressing RFP (no system) or Ec1ApeA. Data show measured phage titer t*_n_* divided by that measured immediately after infection t*_0_*. Cells were infected at an MOI of 0.01. Bars represent the average of three biological replicates with individual data points overlaid. **D,** Top, agarose northern blot of total RNA extracted from *E. coli* expressing RFP (no system), Ec1ApeA, or mutant Ec1ApeA (R521A), infected with FrSangria at an MOI of 10. The blot was probed for the 5’ end of the phage genome. Asterisk, full-length genome of FrSangria. Bottom, the same agarose gel stained with GelRed as loading control before transfer. Data are representative of three biological replicates. p.i., post infection.

To further investigate how ApeA restricts phage replication, we performed time-lapse microscopy to track high MOI infection of single cells expressing either Ec1ApeA or GFP (green fluorescent protein) under the native Ec1ApeA promoter that were mixed at a 1:1 ratio. GFP was used as control instead of RFP for these experiments since the emission spectra of RFP overlap with propidium iodide. In agreement with our liquid infection experiments, the GFP-positive control cells were rapidly lysed by FrSangria within ∼60 min, as was evident by staining with propidium iodide (Figure 2B, Video S1). On the contrary, the majority of cells expressing Ec1ApeA were not lysed by the phage but instead stayed intact and were dividing, indicating that they had successfully thwarted the infection (Figure 2B, Video S1). We then measured the replication of FrSangria by tracking the amount of newly emerging phages from infected cultures of *E. coli* expressing either wild-type Ec1ApeA or RFP. This showed a substantial decrease in phage replication (>400-fold reduction in emerging infectious phages), agreeing with our results from plaque assays, liquid infections, and microscopy (Figure 2C).

Having established that Ec1ApeA does not induce cell lysis, we focused on the identification of the RNA target that is cleaved by the HEPN domain of Ec1ApeA. Based on our liquid infection and microscopy experiments, we did not expect Ec1ApeA to cleave essential cellular RNAs to induce cell death or growth arrest, as it would typically be the case for effector RNases of abortive infection systems^52^. Furthermore, Ec1ApeA did not appear to downregulate expression of the F-pilus, which is the receptor of the *Qubevirus* and *Emesvirus* families of phages in our panel^40,53^, as Ec1ApeA did not influence infection by phages of the *Emesvirus* family (Figure 1B, C). We therefore hypothesized that Ec1ApeA might instead cleave the genome of the infecting RNA phage. To test this hypothesis, we infected bacteria expressing Ec1ApeA, the Ec1ApeA HEPN mutant (R521A), as well as control cells with FrSangria at high MOI and isolated total RNA at several timepoints post infection. We then performed agarose northern blotting of the isolated RNA with a radiolabeled probe specific to the 5’ end of the phage genome (Figure 2D). In bacteria expressing Ec1ApeA, substantial amounts of genome cleavage products (∼200–1200 nt) accumulated from 45 min post infection onward. Crucially, these cleavage products were not apparent when the bacteria expressed the mutated Ec1ApeA protein, indicating that Ec1ApeA indeed cleaves the phage genome in a HEPN-dependent fashion (Figure 2D). In the case of Ec1ApeA-mediated defense against DNA phages, it was recently proposed that it induces cell death via tRNA cleavage^39^. In the case of infection with RNA phages, we did not see evidence for induced cell death by Ec1ApeA (Figure 2A, B), and we observed only minimal tRNA cleavage, suggesting phage genome cleavage to be the main driver of defense (Figure S2B). Together, these results show that ApeA stops RNA phage replication by directly cleaving the phage genome.

### A conserved pocket in the ApeA protein senses RNA phage infection

An important step in antiphage defense is the sensing of the infection by the defense system. To understand how ApeA detects the presence of RNA phages within the cell, we first inspected the AlphaFold 3 structural prediction of Ec1ApeA (Figure 1E) to search for a potential sensor structure. We identified a deep, positively charged pocket on either side of the Ec1ApeA dimer, distal to the dimer interface that forms the HEPN active site (Figure 3A, B). To analyze whether this pocket is a conserved structural feature of ApeA proteins, we first used an AlphaFold 3 prediction of the Ec1ApeA monomer as query to search for structural homologs using Foldseek. From the 362 homologs detected in the AlphaFold50 database^54^, structures with a probability score of >0.5 (248 in total, Table S1) were then used as input in FoldMason^55^ to generate a multiple structural alignment, resulting in an LDDT (local distance difference test) score of 0.314 for the alignment. This analysis revealed that the pocket is a conserved structural feature of ApeA, even though the amino acid identity among the homologs identified by Foldseek was as low as 6% (Figure 3C). In addition to the identified structural conservation of the pocket, ConSurf^56^ analysis identified several highly conserved residues (E28, T34, and R485) positioned at the bottom of the pocket that appeared solvent-exposed and which therefore are likely to be functionally important (Figure 3B, S3A).

**Figure 3.**
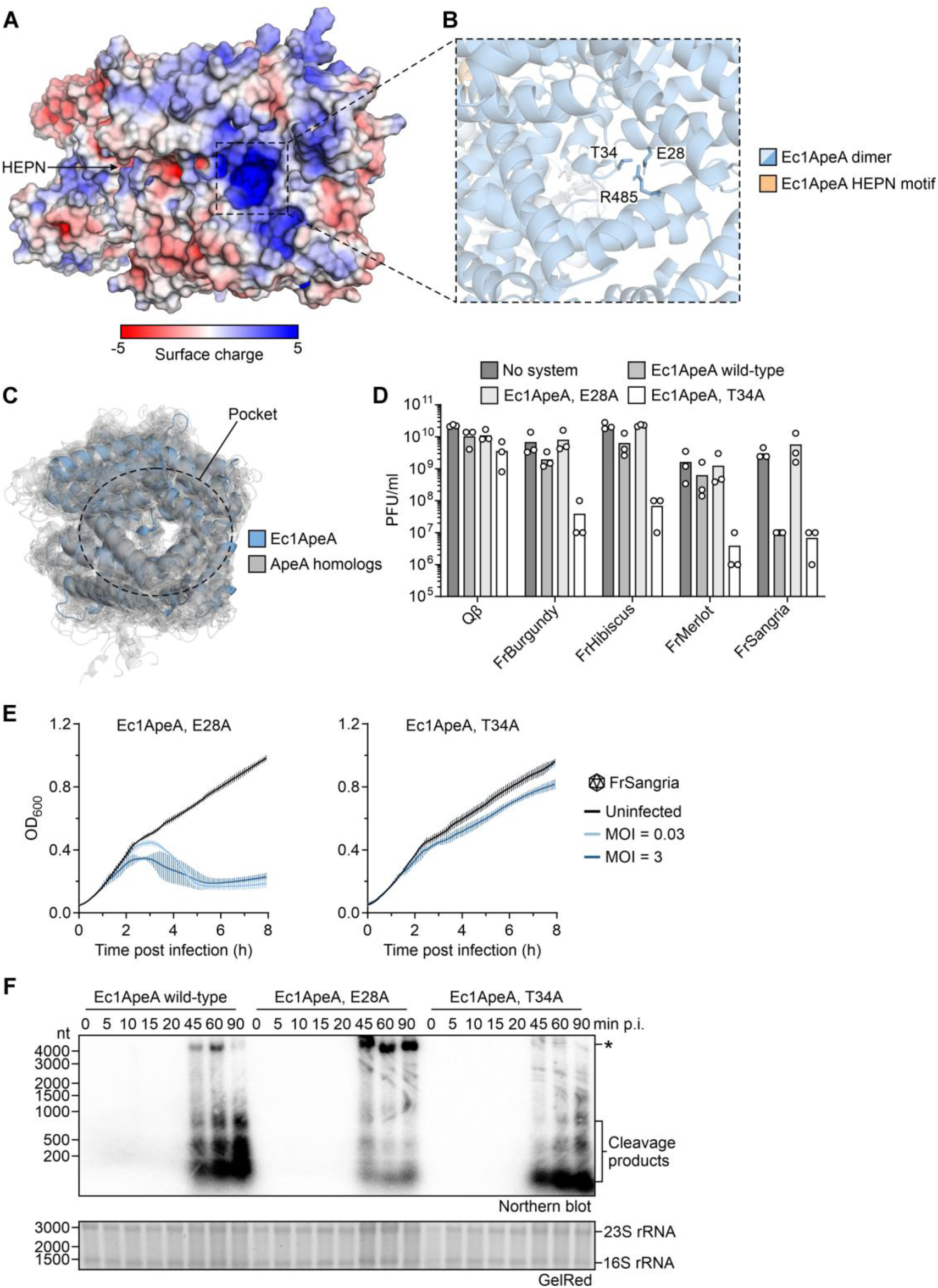
A conserved pocket in the ApeA protein is involved in phage recognition. **A,** Predicted structure of the Ec1ApeA homodimer (compare to Fig. 1E). Surface charge was calculated using the APBS Electrostatics plugin in PyMOL^58^ v3.1.6.1. A positively charged pocket (dashed box) was detected. **B,** Zoom-in of the positively charged pocket shown in **A**. E28, T34, and R485 are fully conserved solvent-exposed residues inside of the pocket that are in close proximity to each other. **C,** Homologs of Ec1ApeA identified by Foldseek^44^, aligned to the predicted structure of the Ec1ApeA monomer using FoldMason^55^. All of the homologs comprise the identified pocket. For clarity, only the top 100 Foldseek hits are shown. **D,** PFU quantification of *Qubevirus* phages infecting *E. coli* expressing pocket mutants of Ec1ApeA (E28A, T34A). Since these experiments were performed together with the plaque assays shown in Fig. 1D, the no system control as well as wild-type Ec1ApeA are repeated for easier comparison. Bars represent the average of three biological replicates with individual data points overlaid. **E,** Growth curves of *E. coli* expressing pocket mutants of Ec1ApeA (E28A, T34A), either uninfected or infected with FrSangria at an MOI of 0.03 or 3. The average with s.d. of three biological replicates is shown. **F,** Top, agarose northern blot of total RNA extracted from *E. coli* Ec1ApeA, or Ec1ApeA mutants (E28A, T34A), infected with FrSangria at an MOI of 10. The blot was probed for the 5’ end of the phage genome. Asterisk, full-length genome of FrSangria. Bottom, the same agarose gel stained with GelRed as loading control before transfer. Data are representative of three biological replicates. p.i., post infection.

To test if the pocket is involved in ApeA-mediated defense, the conserved amino acids identified by ConSurf were substituted with alanine, the constructs transformed and the resulting strains infected with our panel of RNA phages. The R485A substitution showcased strong toxicity, thus preventing us from testing its defensive properties (Figure S3B). While the E28A substitution fully abolished defense against FrSangria, the T34A substitution had no negative influence on the defensive properties of Ec1ApeA (Figure 3D, S3C). On the contrary, while wild-type Ec1ApeA only showed weak defense against the *Qubevirus* phages closely related to FrSangria (Figure 1B), the T34A substitution resulted in a substantial decrease in PFUs for these phages compared to the control (Figure 3D). Liquid infection assays confirmed that the E28A substitution eliminated defense, while the T34A substitution increased defense compared to wild-type Ec1ApeA, indicating that the pocket is involved in RNA phage detection (Figure 3E). Furthermore, we did not observe degradation of the phage genome for the E28A substitution even though this protein had an intact HEPN domain, whereas the T34A substitution strongly inhibited phage replication, as indicated by very weak signals for the full-length phage genome (Figure 3F).

To our surprise, the E28A substitution had varying effects on the protection against DNA phages: No change in defense was observed for T2, while it was fully abolished for T4 (Figure S3D). For T5, the E28A substitution showed less defense than wild-type Ec1ApeA in plaque assays, but wild-type protection in liquid infection (Figure S2A, S3D). Another homolog of ApeA, Ec2ApeA, was recently shown to sense DNA phage infection via binding of deoxydinucleotides to the pocket, which were predicted to accumulate during phage-induced host genome degradation^39^. Since the RNA phages we employed in our experiments do not degrade the host genome^57^, it is unlikely that accumulation of deoxydinucleotides triggers Ec1ApeA activity in RNA phage defense. Indeed, deoxydinucleotides do not activate Ec1ApeA^39^. These observations provide further evidence that the mechanism of defense of ApeA differs between RNA and DNA phages. Together, these results show that ApeA contains a conserved positively charged pocket that is involved in the detection of an ongoing infection and which is essential for defense against RNA phages.

### An RNA structure in the phage genome activates ApeA

Next, we investigated which factor of the invading RNA phage is sensed by the conserved pocket of Ec1ApeA. To that end, we isolated escaper mutants of FrSangria that are able to overcome the defense conferred by Ec1ApeA. We first picked 20 single plaques that formed on double-layer agar plates of bacteria expressing Ec1ApeA infected with parental FrSangria. These 20 potential escaper mutants were then propagated in liquid culture on bacteria expressing Ec1ApeA. This procedure was repeated once to obtain the final lysates of potential escaper mutants. The obtained phages were then used in plaque assays to infect cells expressing Ec1ApeA and compared to an RFP-expressing control strain. Of the 20 isolated phages, 11 robustly escaped defense by Ec1ApeA (although Ec1ApeA still reduced the plaque size of most of them), indicating that they had acquired escaper mutations (Figure 4A). To further verify the escape, we performed time-lapse microscopy of bacteria infected with one escaper mutant (Esc-20) as we did before for wild-type FrSangria (Figure 2B). This mutant was able to synchronously lyse both GFP-expressing control cells and cells expressing Ec1ApeA, demonstrating that it can successfully escape defense by Ec1ApeA (Figure 4B, Video S2).

**Figure 4.**
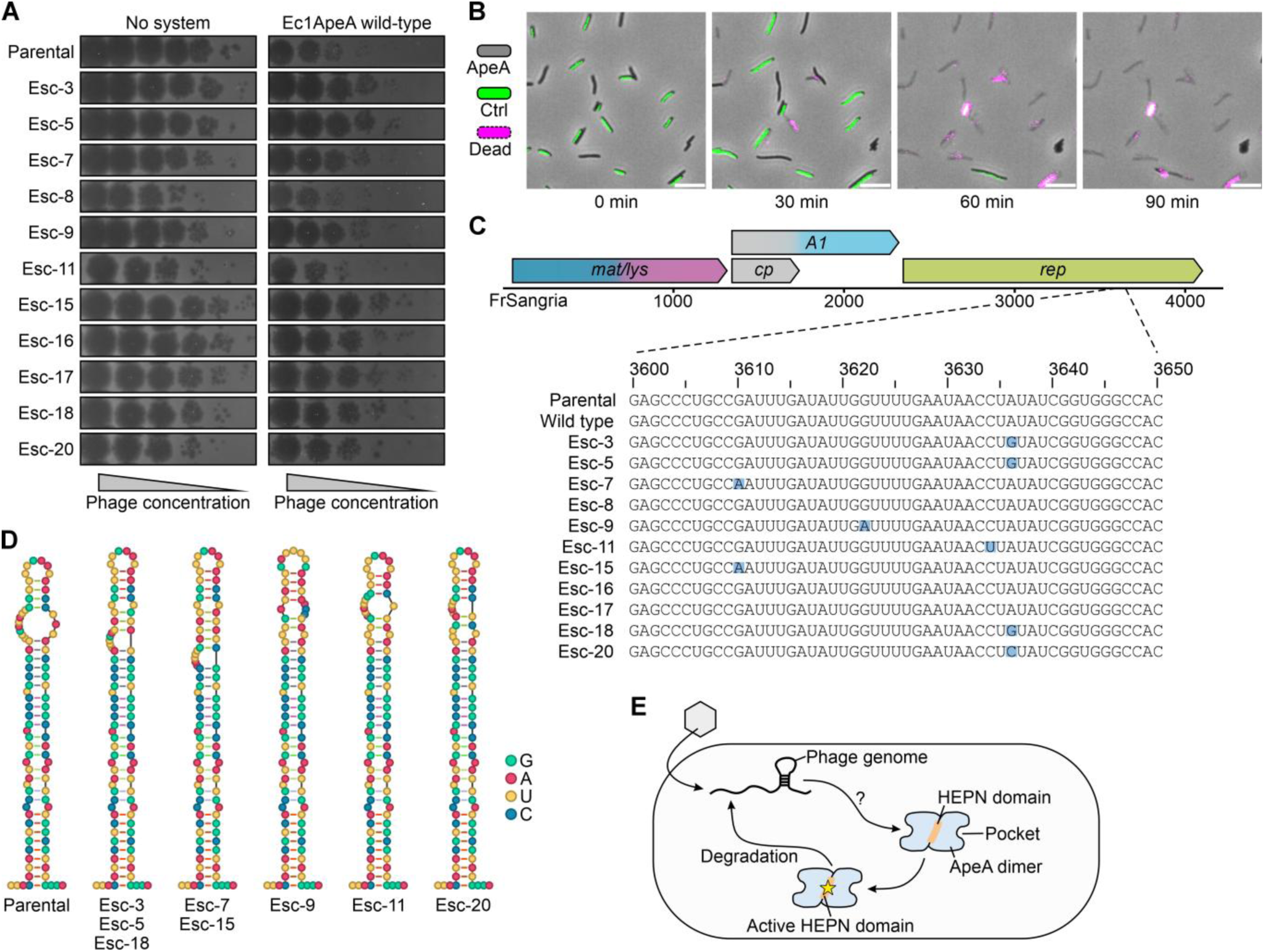
FrSangria can escape ApeA defense by mutating a stem loop in its RNA genome. **A,** Plaque assays of the parental lysate of FrSangria used to generate phage mutants and the resulting escaper phages (Esc-3 to Esc-20) on lawns of *E. coli* expressing RFP (no system) or wild-type Ec1ApeA. All escaper mutants plaque similarly on Ec1ApeA and the control, albeit with smaller plaques. **B,** Time-lapse microscopy of live *E. coli* cells expressing GFP (green) mixed with cells expressing Ec1ApeA (black). Cells were infected with Esc-20 at an MOI of 5 in the presence of propidium iodide (PI, magenta) and incubated at 37°C in an agar pad. The three channels (phase contrast, GFP, PI) are presented as merged images. Scale bar, 10 µm. Data are representative of three biological replicates. **C,** Top, genome organization of FrSangria (4215 nt, NCBI ID: PP430145.1). mat/lys, maturation protein with additional lysis protein function. cp, coat protein. A1, minor capsid protein that is a translational read-through product of *cp*. rep, replicase protein. Bottom, multiple sequence alignment of positions 3600–3650 of the genome of FrSangria comparing the sequences of the parental phage, the escaper phages, as well as an additional wild-type lysate that was not used for escaper generation. Blue, mutated nucleotides in the escaper phages. **D,** Secondary structures of positions 3579–3668 of the genomic RNA of FrSangria, predicted using RNAfold^59^. The centroid structures are presented. **E,** Model of the mechanism of action of ApeA. Following infection with an RNA phage, a structure in the genomic RNA of the phage activates the HEPN domain of ApeA, likely by interacting with a conserved pocket. This induces degradation of the phage genome to stop the infection.

To assess the mutations the escaper phages had acquired, we isolated the genomes of the 11 escaper phages (Figure 4A), reverse transcribed them, and subjected them to nanopore sequencing. Comparison of the genomic sequences of the escaper phages to the parental phage as well as an additional wild-type genome revealed that all escaper phages had acquired several mutations (Tables S2, S3). Around half (31/60) of these mutations were silent and are therefore not expected to influence the function of the respective phage proteins. Crucially, 8 of the 11 escaper phages had mutations in a short sequence located between positions 3600–3636 of the genome (within the replicase gene) that were either silent or introduced amino acid substitutions that are expected to only have a small influence on the protein (V to I or D to N) (Figure 4C, Table S3). This region therefore emerged as a hotspot for mutations that allow escape from ApeA defense.

It is well-known that the genomes of RNA phages are strongly structured^17^. We therefore hypothesized that the mutations identified in this hotspot of the escaper phages influenced an RNA structure in the genome that triggers ApeA. To test this hypothesis, we predicted the secondary RNA structures around the mutation hotspot using RNAfold^59^. These predictions showed that the wild-type genomic RNA forms a long stem loop between positions 3582–3665 that has a bulge close to its apical loop (Figure 4D). This stem loop was previously shown to be in contact with coat protein dimers within mature virions of Qβ, although the functional relevance of this remains unclear^60^. The mutations identified in the escaper phages were predicted to reduce the size and/or position of the bulge of the stem loop by restructuring the apical part of the stem (Figure 4D). This indicates that structural changes in a stem loop in the genomic RNA of FrSangria might allow escape from ApeA defense. Consistently, the genome of one of the escaper phages (Esc-3) exclusively harbored silent mutations (Table S3), suggesting that this phage might indeed escape based solely on RNA structure changes. We then predicted the structures of the same stem loop in the genomes of the other *Qubevirus* phages we used in our study (Figure S4). In Qβ, the apical bulge of FrSangria is absent, whereas the basal half of the stem is more open than that of FrSangria (Figure S4). In the genomes of FrBurgundy, FrHibiscus, and FrMerlot, the stem loop has the same apical structure as the one of FrSangria, whereas the basal structure is like the one in Qβ (Figure S4). These observations could explain why Ec1ApeA does not defend against Qβ, while it causes a clear reduction in plaque size for FrBurgundy, FrHibiscus, FrMerlot (Figure 1B). Together, the results of our escaper assay suggest that ApeA recognizes an RNA structure in the genome of RNA phages, which then triggers phage genome degradation to stop the infection (Figure 4E).

## Discussion

Over the last years, an astonishing diversity of bacterial antiphage defense systems has been discovered. Yet we know surprisingly little about the mechanisms of defense against infection by RNA phages, even though recent studies show that these phages are much more abundant and diverse than previously anticipated^14–16^. In this study, we investigated RNA phage defense conferred by ApeA and found that Ec1ApeA strongly protects against the Qβ-like RNA phage FrSangria. Our data suggest that Ec1ApeA senses an RNA structure within the phage genome, which triggers the activation of its HEPN RNase domain to cleave the phage genomic RNA to ultimately stop the infection (Figure 4E).

Although we identify a conserved pocket within Ec1ApeA and confirm that it is essential for defense against RNA phages, the exact mechanism of how it is involved in the sensing of the invading phage remains to be studied. Our results suggest that Ec1ApeA might recognize an RNA structure in the phage genomic RNA, as we identified a structured region that was repeatedly mutated in escaper phages. Whether these mutations only cause a change in secondary structure or if longer range tertiary structures that could be sensed by Ec1ApeA are also affected will be examined in future work. In addition to its pocket, Ec1ApeA has a strongly positively charged face that is on the opposite side of the HEPN active site (Figure S5). This positively charged face is reminiscent of other RNA-binding proteins such as Hfq^61^, and could be involved in the binding or coordination of the phage genomic RNA to make it accessible to the HEPN domain. Interestingly, different homologs of ApeA seem to sense different factors to become activated, which further depends on the type of phage that is infecting the cell. During DNA phage infection, Ec2ApeA is activated by deoxydinucleotides, while Ec1ApeA is not^39^. How Ec1ApeA is activated by DNA phages remains unknown, but one hypothesis is that it might sense structured RNAs similar to what we suggest for RNA phages. Activation of defense systems by structured RNAs expressed by DNA phages has previously been shown for CBASS (cyclic oligonucleotide-based antiphage signaling system)^62^, indicating that RNA sensors may be important in phage defense, similar to eukaryotic antiviral immunity^13^.

A striking characteristic of RNA phage defense by Ec1ApeA is that it directly cleaves phage genomic RNA, similar to restriction modification^46^ or CRISPR-Cas systems^63^. This cleavage is dependent on both the HEPN RNase domain as well as the pocket, as disruption of either inhibits genome cleavage. We also observe minor cleavage of host tRNAs upon RNA phage infection, which was proposed to be the main function of ApeA in DNA phage defense^39^. Yet RNA phage infection does not induce ApeA-mediated cell death, demonstrating that direct cleavage of the phage genome is the main mode of action of ApeA for these phages. These observations suggest that there is a substantial difference between ApeA-mediated defense against DNA and RNA phages. Phage genome cleavage by ApeA is also different from the targets of other HEPN domain-containing antiphage systems like type III CRISPR-Cas systems or toxin-antitoxin system that typically induce cell death/growth arrest via degradation of host RNAs^21,50,64,65^. Our results thus add to the growing repertoire of targets of antiviral HEPN domain-containing protein, further expanding the scope of this widespread domain that is conserved from bacteria to mammals^29,30^.

While our study primarily focused on a single model phage and how its replication is inhibited by Ec1ApeA, we also identified a second homolog of ApeA, Ps2ApeA, which is able to broadly restrict RNA phages of the *Fiersviridae* family. Although Ps2ApeA is only ∼31% identical to Ec1ApeA at the amino acid level, it is placed in the same phylogenetic clade as Ec1ApeA. In contrast to Ps2ApeA, other ApeA homologs we tested here do not protect against RNA phage infection. It is therefore possible that the Ec1/Ps2ApeA clade of ApeA proteins has evolved to defend against both RNA and DNA phages. Further testing of a wider range of ApeA homologs would be required to corroborate this hypothesis.

In summary, we have revealed that ApeA, which is a widespread defense system consisting of only one protein, directly cleaves the genome of an infecting RNA phage to restrict its replication. Activity of ApeA does not induce cell death and therefore stands out as a non-abortive defense system. Activation of ApeA is likely triggered by an RNA structure in the phage genome, which may occur immediately upon genome injection or later during genome replication. Our findings provide important insights into the poorly understood mechanisms of bacterial defense against RNA phages.

## Materials and methods

### Bacterial strains and phages

For all experiments with F-dependent RNA phages or DNA phages, the ApeA system (NCBI ID: WP_000706972.1) and its mutant variants were expressed in *E. coli* W1485, which is F+. In case of infection with phage C-1, *E. coli* JE-1, which harbors an IncC plasmid, was used. As a negative control for all experiments, a plasmid carrying either *rfp* or *gfp* in place of the *apeA* gene was used (referred to as “no system”). Cultures were grown in MMB (LB supplemented with 5 mM MgCl₂ and 0.1 mM MnCl₂) at 37°C with shaking at 200 rpm, with the appropriate antibiotics added (20 µg/ml chloramphenicol or 10 µg/ml gentamicin). The bacterial strains and phages used in this study are listed in Table S4. Infections were carried out in MMB, either with or without 0.5% agar.

### Phage storage and lysate preparation

For long-term storage of RNA phages, the host strain at exponential phase was infected with the respective phage at an MOI of ∼1 and incubated for 30 min at 37°C with shaking at 200 rpm. Glycerol was added to the infected bacteria to a final concentration of 25% and the stocks stored at −80 °C. To prepare phage lysates, an overnight culture of the host strain was diluted 1:1000 in MMB and infected with a scrape of frozen phage stock. The infected culture was incubated overnight at 37 °C with shaking at 200 rpm. The following day, cultures were centrifuged for 2 min at 13,000 rcf at RT, and the supernatant was filtered through a 0.2 µm filter to remove remaining bacteria and cellular debris. The resulting lysates were stored at 4 °C and their titers determined using plaque assays.

### Plasmid and strain construction

All plasmids and their corresponding inserts used in this study are listed in Table S5. *apeA* variants were generated by amplifying the plasmid containing wild-type *apeA* using primers that included the intended modification, followed by treatment with KLD enzyme mix (NEB) according to the manufacturer’s instructions to generate transformable plasmids. Alternatively, PCR reactions were treated with DpnI (NEB) and assembled using TEDA (T5 exonuclease DNA assembly)^68^. After sequence verification, the plasmids were transformed into *E. coli* W1485 using TSS (transformation and storage solution)^69^.

### Plaque assays

The defensive activity of the ApeA system and its mutant variants was assessed using the small drop plaque assay^70^. For each condition, 300 μl of an overnight bacterial culture carrying the relevant ApeA construct (or the negative control strain) was combined with 30 ml of molten 0.5% MMB agar. This mixture was poured into a 12 cm square plate and allowed to solidify for 15 min at room temperature. For IPTG-inducible plasmids, 100 µM IPTG was added to the agar prior to pouring, and plates were then incubated for 1 h at room temperature. Phages were serially diluted ten-fold in MMB, and 5 or 10 μl drops of each dilution were applied onto the bacterial lawn. Plates were incubated overnight at 37°C, and plaque-forming units (PFUs) were counted after overnight incubation. In case only a faint zone of lysis without individually discernible plaques was visible, this was counted as 10 plaques.

### Liquid infection assays

Single colonies of each strain were grown in MMB at 37°C with shaking at 200 rpm to late exponential phase and diluted at least 1:100 in MMB to obtain the same OD_600_ for each strain. The cultures were then grown until reaching an OD_600_ of 0.2, after which 180 μl of culture was transferred into a 96-well plate. Next, 20 μl of either MMB (uninfected control) or phage lysate was added to achieve a final MOI of 3 or 0.03. Plates were incubated at 37°C with shaking in a Tecan Infinite 200 Pro plate reader, and OD_600_ measurements were taken every 5 min. Infections were conducted as biological triplicates across three separate plates, with technical triplicates within each plate. In each plate, three wells containing only medium served as blanks, and the averages of these values were subtracted from the OD_600_ readings of wells containing bacteria.

### Time-lapse microscopy

An overnight bacterial culture was diluted 1:1000 and grown at 37°C with shaking at 200 rpm to an OD_600_ of 0.2. Cells were infected with phages at an MOI of 5 for 15 min at 37 °C in a total volume of 300 µl to allow adsorption. Following adsorption, cells were pelleted by centrifugation at 5000 rcf for 5 min, the supernatant discarded, and the pellet resuspended in 75 µl of 1x PBS supplemented with 0.2 µg/ml propidium iodide (PI). Cells were then placed on a 1% agarose pad prepared in 20% MMB in 1x PBS. Time-lapse imaging was performed using an epifluorescence microscope (Leica) equipped with an LED light engine for fluorescence illumination and a DFC9000GT-VSC11903 camera. Cells were maintained at 37 °C using an Incubator i8 (PECON) during imaging. Images were acquired every 3 min for 90 min using a 40x air objective.

### Phage replication assay

Overnight cultures of strains expressing either ApeA or RFP were diluted 1:1000 in MMB and grown at 37°C with shaking at 200 rpm to an OD_600_ of 0.2. The cultures were infected with FrSangria at an MOI of 0.001. For each timepoint, 1 ml of the infected culture was collected, cells were pelleted by centrifugation for 2 min at 13,000 rcf at RT, and 10 µl of the supernatant was used to prepare 10-fold serial dilutions. Plaque assays for the supernatant of each timepoint were performed as described above with *E. coli* W1485 as the indicator strain.

### Growth curve assays

Overnight cultures were diluted 1:100 in a 96-well plate and incubated at 37°C with shaking in a Tecan Infinite 200 Pro plate reader, with OD_600_ measurements taken every 10 min. Three biological replicates with technical triplicates for each strain were performed. Three wells containing only medium served as blanks, and the average of these values were subtracted from the OD_600_ readings of wells containing bacteria.

### RNA extraction

For RNA extraction, bacteria expressing RFP, ApeA, or ApeA mutants were grown to an OD_600_ of 0.2 at 37°C with shaking at 200 rpm. 2 ml of the culture was collected as the uninfected control (t = 0 min) before infecting the remaining culture with FrSangria at an MOI of 10. 2 ml of the infected samples were collected for each timepoint. Directly after collection, the samples were centrifuged at 4°C and 7,000 rcf for 1 min to pellet the bacteria, the supernatant was discarded und the pellets snap-frozen in liquid nitrogen before being stored at −80°C. The frozen pellets were then thawed on ice and total RNA was extracted with 1 ml of TRIzol (Invitrogen) according to the manufacturer’s protocol. To get rid of contaminating DNA, 0.5 µl of water, 5 µl DNase I buffer with MgCl_2_ (Thermo Scientific), 0.5 µl of RNase inhibitor (Thermo Scientific) and 4 µl DNase I (Thermo Scientific) was added and the mixture incubated at 37°C for 30 min. The DNase-digested RNA was subjected to acidic phenol-chloroform extraction and finally dissolved in 30 µl of water.

### Agarose northern blotting

1 µg of total RNA was mixed with RNA loading buffer, denatured for 2 min at 70°C and put on ice. The RNA was then run in a denaturing 1% TBE agarose gel containing 0.1% sodium hypochlorite (Carl Roth). As loading control, the gel was stained in a bath containing GelRed (Sigma-Aldrich) at 1:10,000 dilution. The RNA was transferred to a nylon membrane (Sigma Aldrich) by capillary transfer overnight in the presence of 10x SSC. Membranes were UV-crosslinked at 254 nm and hybridized with a radiolabeled DNA oligonucleotide (GGTGTGTGTCGGAAGATTCGA) specific to the 5’ end of the FrSangria genome at 42°C in Roti-Hybri-Quick hybridization buffer (Carl Roth). Membranes were washed in three subsequent steps with 5x SSC, 1x SSC, and 0.5x SSC, all containing 0.1% SDS. Finally, the membrane was dried, exposed to a phosphor screen, and imaged using a FLA-3000 phosphorimager (Fujifilm).

### RNA PAGE

1 µg of total RNA was mixed with RNA loading buffer, denatured for 2 min at 70°C and put on ice. The RNA was then separated in a 6% polyacrylamide gel containing 7 M urea in 1x TBE for 2 h at 300 V. For visualization of the RNA, the gel was stained for 15 min in a bath containing SYBR Green I (Invitrogen) in 1x TBE at 1:10,000 dilution before being imaged using a FLA-3000 phosphorimager.

### Polyacrylamide northern blotting

1 µg of total RNA was mixed with RNA loading buffer, denatured for 2 min at 70°C and put on ice. The RNA was then separated in a 6% polyacrylamide gel containing 7 M urea in 1x TBE for 2 h at 300 V. The RNA was transferred to a nylon membrane for 1 hour at 50 V in 1x TBE. The RNA was UV-crosslinked to the membrane at 254 nm and hybridized overnight at 42°C in Roti-Hybri-Quick hybridization buffer with a radiolabeled DNA oligonucleotide (GATTCGAACCCCCGATACG) specific to the 3’ end of tRNA-Ser^GGA^. The membrane was washed and imaged as described above.

### Phylogenetic analysis of ApeA

To retrieve homologs of ApeA, the originally published ApeA from *E. coli* NCTC8008 (Ec1ApeA)^7^ was used as the query to perform three iterations of PSI-BLAST^43^ searches against the non-redundant database of NCBI using default parameters, and the resulting 500 sequences were downloaded. In addition, an AlphaFold 3^42^ structural prediction of ApeA was used to query the AlphaFold50 database^54^ using Foldseek^44^ in TM-align mode. Of the 362 homologs detected, structures with a probability score of >0.5 (248 in total) were added to the 500 sequences obtained via PSI-BLAST. The combined list of 748 proteins (Table S1) was clustered using MMseqs2^71^ with a minimum sequence identity of 0.9 and a minimum alignment coverage of 0.8 in normal mode. The final list was aligned with Clustal Omega^72^ using default parameters, followed by clipping using ClipKIT^73^ v2.10.2 with -m smart-gap. A phylogenetic tree was generated using IQ-TREE^74^ v3.0.1 with -m Q.pfam+F+R10 -bb 1000 -alrt 1000 and finally visualized using iTOL^75^ v7.

### Structural predictions

Structural prediction of ApeA was performed using AlphaFold 3. Visualization as well as calculation of the surface charge via the ABPS Electrostatics plugin was performed in PyMOL^58^ v3.1.6.1. Analysis of conserved residues was performed using ConSurf^56^ with the AlphaFold 3 prediction of ApeA and the multiple sequence alignment of the 500 ApeA homologs identified via PSI-BLAST as the inputs. Multiple structural alignment was performed using FoldMason^55^ in easy-msa mode with --report-mode 1 and the Foldseek hits described above (Table S1) as inputs.

### Generation of escaper phages

To generate escaper phages, a wild-type lysate of FrSangria (referred to as “parental”) was plated on double-layer agar plates using *E. coli* expressing ApeA. Briefly, 150 µl of an overnight bacterial culture was mixed with 15 ml of molten 0.5% MMB agar. The mixture was supplemented with parental phage lysate diluted to yield ∼100 PFU in the plate and poured on top of a petri dish with 1.5% MMB agar. The plate was incubated overnight at 37 °C. Single plaques that formed on this plate were picked into 100 µl of MMB medium. Next, 2 µl of this suspension was used to propagate the selected phages overnight on *E. coli* expressing ApeA. The following day, infected cultures were centrifuged for 2 min at 13,000 rcf at RT, and the supernatant was transferred to a new tube containing 1-2 drops of chloroform to lyse any remaining bacteria. Plaque assays were performed to test the ability of the propagated phages to escape ApeA defense by comparing plaquing on bacteria expressing either RFP or ApeA. For each escaper candidate, a single plaque from the plate with the ApeA-expressing strain was again picked into 100 µl of MMB medium, and the selection-propagation procedure was repeated once to obtain the final escaper candidates.

### Sequencing of phage genomes

To extract the genomic RNA of FrSangria and escaper phages, 200 µl of the phage lysates (∼10^10^ PFU) were supplemented with 1 µl of DNase I and 22 µl of 10x DNase I reaction buffer and incubated for 30 min at 37°C. Then, the genomic RNA was extracted in 1 ml of TRIzol (Invitrogen) according to the manufacturer’s protocol. 1 µg of purified phage genome was polyadenylated with *E. coli* poly-A polymerase (Invitrogen) following the manufacturer’s protocol. Polyadenylated RNA was purified using the RNA Clean & Concentrator-5 kit (Zymo Research) according to the manufacturer’s protocol. To generate cDNA and finally dsDNA, the UltraMarathonRT Template Switching Kit (RNAConnect) was used according to the manufacturer’s protocol. The final products were cleaned up using a column-based kit (Macherey-Nagel) and subjected to nanopore sequencing (Plasmidsaurus).

To generate a reference for the identification of mutations in the escaper phage genomes, a consensus sequence of the parental phage was first generated. To that end, the nanopore sequencing data of the parental phage was used as input in medaka_consensus v2.2.0 (https://github.com/nanoporetech/medaka) and compared against the FrSangria NCBI reference genome (PP430145.1) with -m r1041_e82_400bps_sup_v5.2.0. The generated parental consensus sequence was then used to call variants in the escaper phages using medaka_variant v2.2.0 (https://github.com/nanoporetech/medaka) with -m r1041_e82_400bps_sup_variant_v5.0.0. Consensus sequences for each escaper phage were generated using medaka_consensus as above. Additionally, an independent wild-type lysate of FrSangria was sequenced as control for the parental phage and its consensus sequence was generated as for the escaper phages. All genome sequences are listed in Table S2. Finally, a multiple sequence alignment of the parental, the control wild-type, and the escaper phages was generated using Clustal Omega with default parameters to identify mutation hotspots.

## Acknowledgments

We thank Anke Sparmann and Jörg Vogel for comments on earlier versions of the manuscript. We further thank Siân V. Owen at the Wadsworth Center and Michael Baym at Harvard Medical School for generously sharing the Fr RNA phages. Research in the Hör lab is supported by an ERC Starting Grant (grant number 101221218 to J.H.) and the DFG-funded SPP2389 (grant number 569011257 to J.H.).

## Data availability

All data are available in the main text or supplemental information.

## Author contributions

A.D. and J.H. conceived and designed the study. A.D., M.V.G., S.A., and S.R. performed experiments. A.D. and J.H. analyzed the data. J.H. secured funding and supervised the study. J.H. wrote the original draft of the manuscript. All authors reviewed and edited the manuscript and support the conclusions.

---

**Figure S1.**
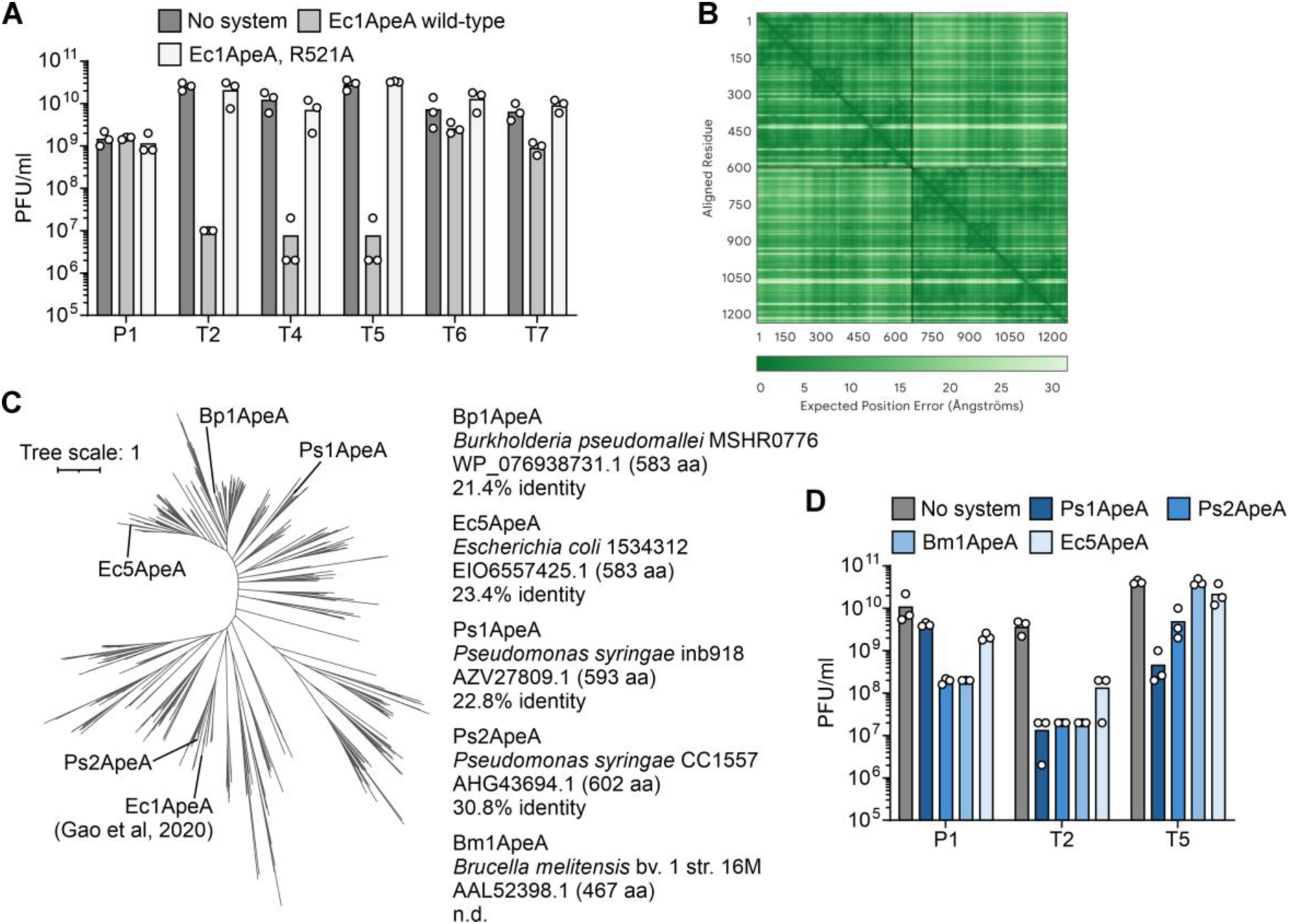
Diversity and defense phenotype of ApeA homologs. **A,** PFU quantification of DNA phages infecting *E. coli* expressing RFP (no system), Ec1ApeA, or the Ec1ApeA R521A mutant. Bars represent the average of three biological replicates with individual data points overlaid. **B,** Predicted alignment error of the AlphaFold 3 prediction shown in Fig. 1E. **C,** Left, unrooted phylogenetic tree of 470 ApeA homologs. Right, ApeA homologs chosen for experimental characterization. For each homolog, the name, its original host, NCBI ID, and amino acid identity compared to Ec1ApeA are presented. BLAST analysis found no sequence similarity for Bm1ApeA (n.d.). **D,** PFU quantification of DNA phages infecting *E. coli* expressing different ApeA homologs. Bars represent the average of three biological replicates with individual data points overlaid.

**Figure S2.**
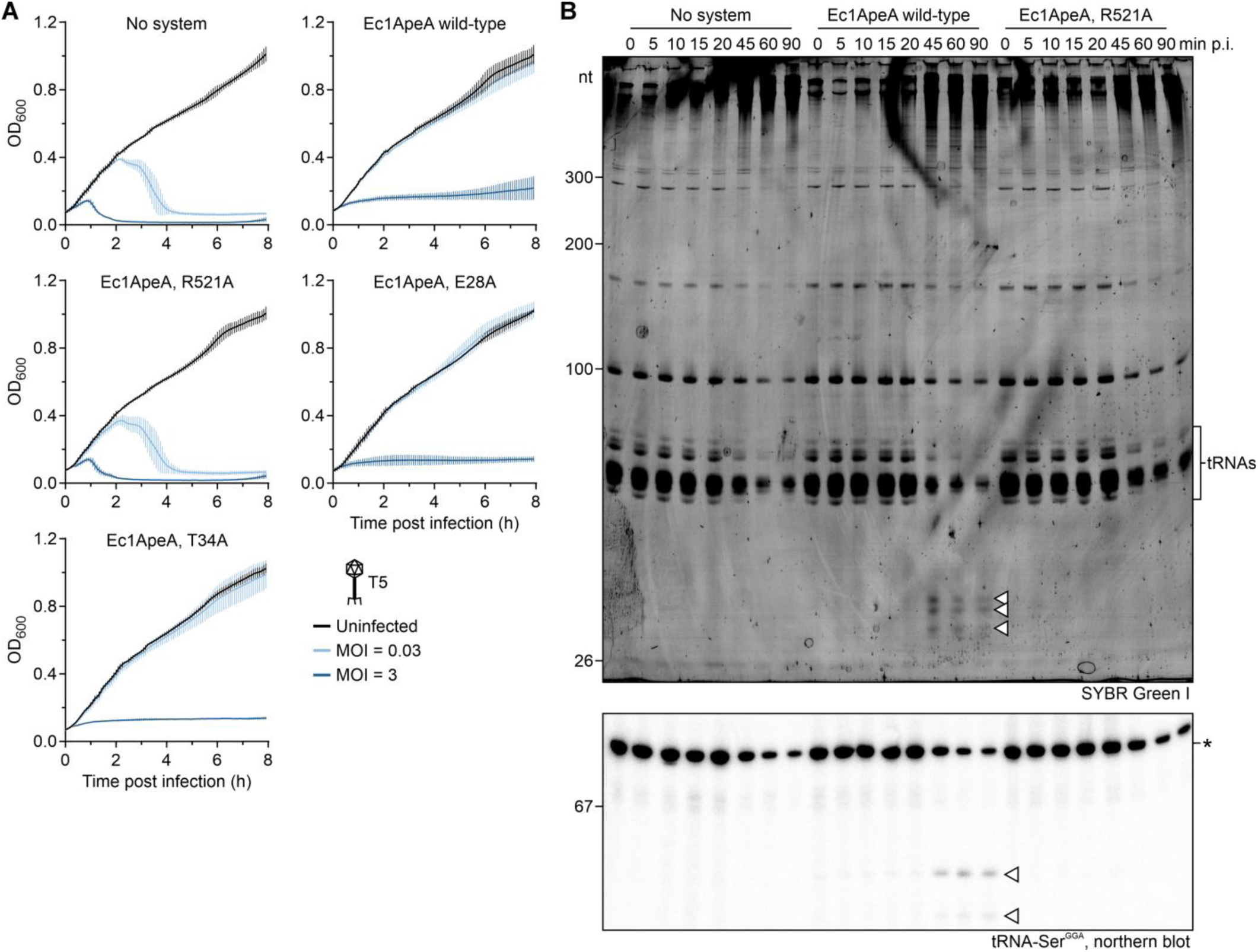
ApeA confers resistance against phage T5 and shows minimal tRNA cleavage upon RNA phage infection. **A,** Growth curves of *E. coli* expressing RFP (no system), Ec1ApeA, or Ec1ApeA mutants (R521A, E28A, or T34A) either uninfected or infected with phage T5 at an MOI of 0.03 or 3. The average with s.d. of three biological replicates is shown. **B,** Top, denaturing urea PAGE of total RNA extracted from *E. coli* expressing RFP (no system), Ec1ApeA, or mutant Ec1ApeA (R521A), infected with FrSangria at an MOI of 10, stained with SYBR Green I. Bottom, northern blot of the same total RNA, probed for the 3’ end of tRNA-Ser^GGA^. Data are representative of three biological replicates. Asterisk, full-length tRNA-Ser^GGA^. White triangles, cleavage products specific to infected cells that express wild-type Ec1ApeA. p.i., post infection.

**Figure S3.**
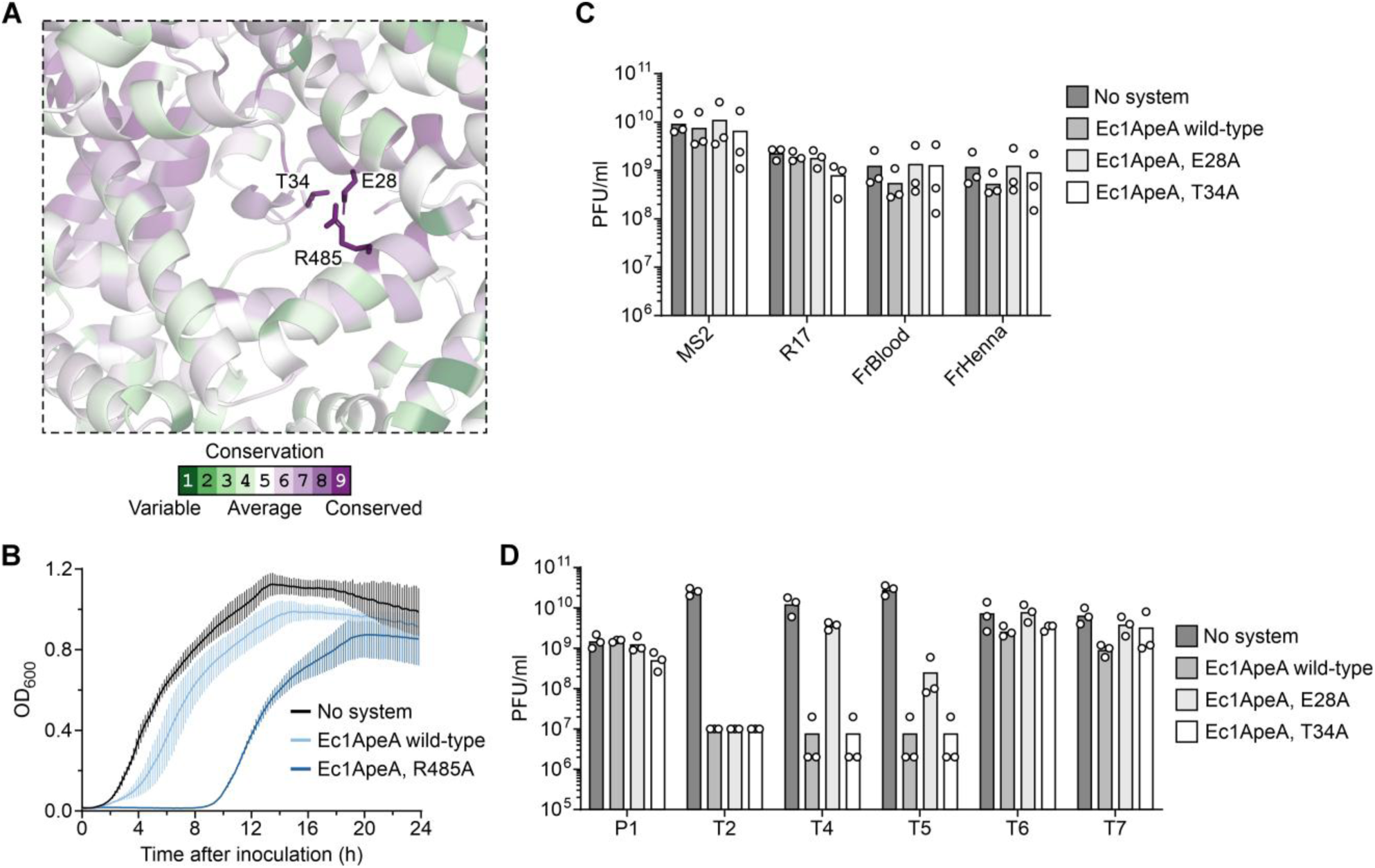
A conserved pocket in the ApeA protein is involved in phage defense. **A,** Conservation analysis of the pocket of ApeA using ConSurf. **B,** Growth curves of *E. coli* expressing RFP (no system), Ec1ApeA, or the Ec1ApeA R485A mutant. Overnight cultures of each strain were diluted 1:1000 and their growth recorded. The average with s.d. of three biological replicates is shown. **C,** PFU quantification of *Emesvirus* phages infecting *E. coli* expressing pocket mutants of Ec1ApeA (E28A, T34A). Since these experiments were performed together with the plaque assays shown in Fig. 1D, the no system control as well as wild-type Ec1ApeA are repeated for easier comparison. Bars represent the average of three biological replicates with individual data points overlaid. **D,** PFU quantification of DNA phages infecting *E. coli* expressing pocket mutants of Ec1ApeA (E28A, T34A). Since these experiments were performed together with the plaque assays shown in **Fig. S1A**, the no system control as well as wild-type Ec1ApeA are repeated for easier comparison. Bars represent the average of three biological replicates with individual data points overlaid.

**Figure S4.**
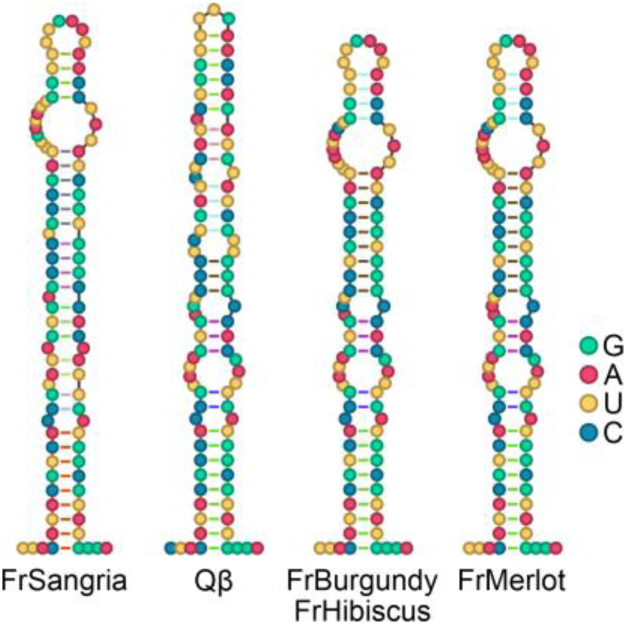
A genomic RNA stem loop structure differs between related *Qubevirus* phages. Secondary structures of a stem loop in the genomic RNA of *Qubevirus* phages, predicted using RNAfold. Genome positions shown are 3579–3668 (FrSangria), 3580–3669 (Qβ), 3586–3675 (FrBurgundy), 3582–3671 (FrHibiscus), and 3586–3675 (FrMerlot). The centroid structures are presented.

**Figure S5.**
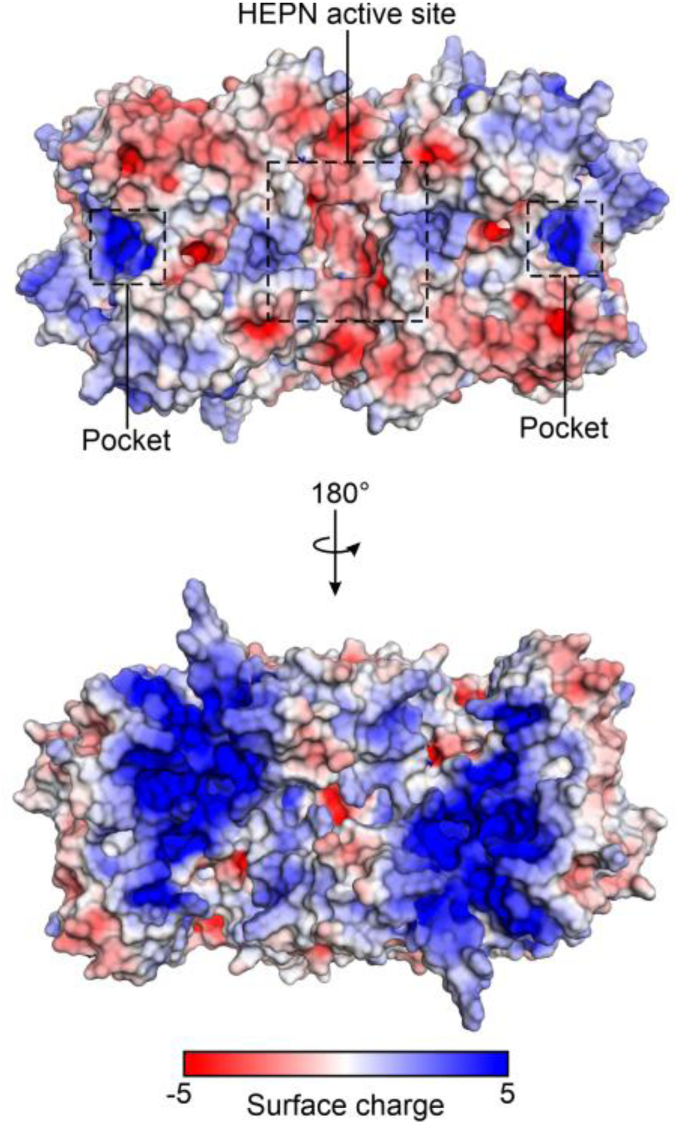
ApeA has a strongly positively charged bac. Predicted structure of the Ec1ApeA homodimer (compare to Fig. 1E). Surface charge was calculated using the APBS Electrostatics plugin in PyMOL v3.1.6.1.

## Legends for supplementary videos

**Video S1. ApeA restricts RNA phage infection.** Time-lapse microscopy of live *E. coli* cells expressing GFP (green) mixed with cells expressing Ec1ApeA (black). Cells were infected with FrSangria at an MOI of 5 in the presence of propidium iodide (PI, magenta) and incubated at 37°C in an agar pad. The three channels (phase contrast, GFP, PI) are presented as a merged video. Scale bar, 10 µm. Data are representative of three biological replicates.

**Video S2. Esc-20 can successfully infect bacteria expressing ApeA.** Time-lapse microscopy of live *E. coli* cells expressing GFP (green) mixed with cells expressing Ec1ApeA (black). Cells were infected with Esc-20 at an MOI of 5 in the presence of propidium iodide (PI, magenta) and incubated at 37°C in an agar pad. The three channels (phase contrast, GFP, PI) are presented as a merged video. Scale bar, 10 µm. Data are representative of three biological replicates.

